# Progesterone shapes medial temporal lobe volume across the human menstrual cycle

**DOI:** 10.1101/2020.02.04.934141

**Authors:** Caitlin M. Taylor, Laura Pritschet, Rosanna Olsen, Evan Layher, Tyler Santander, Scott T. Grafton, Emily G. Jacobs

**Author notes:** Correspondence: Emily G. Jacobs, Dept. of Psychological & Brain Sciences, University of California, Santa Barbara, Santa Barbara, CA 93106, Caitlin M. Taylor, Dept. of Psychological & Brain Sciences, University of California, Santa Barbara, Santa Barbara, CA 93106.

## Abstract

The rhythmic production of sex steroid hormones is a central feature of the mammalian endocrine system. In rodents and nonhuman primates, sex hormones are powerful regulators of hippocampal subfield morphology. However, it remains unknown whether intrinsic fluctuations in sex hormones alter hippocampal morphology in the human brain. In a series of dense-sampling studies, we used high-resolution imaging of the medial temporal lobe (MTL) to determine whether endogenous fluctuations (Study 1) and exogenous manipulation (Study 2) of sex hormones alter MTL volume over time. Across the menstrual cycle, intrinsic fluctuations in progesterone were associated with volumetric changes in CA2/3, entorhinal, perirhinal, and parahippocampal cortex. Chronic progesterone suppression abolished these cycle-dependent effects and led to pronounced volumetric changes in entorhinal cortex and CA2/3 relative to freely cycling conditions. No associations with estradiol were observed. These results establish progesterone’s ability to rapidly and dynamically shape MTL morphology across the human menstrual cycle.

**Highlights:**

- Sex hormones are powerful regulators of hippocampal plasticity in mammals.
- The impact of hormone fluctuations on hippocampal morphology in humans is unknown.
- High resolution imaging of the MTL was conducted across two 30-day periods.
- Progesterone dynamically shapes MTL volume across the human menstrual cycle.
- Chronic progesterone suppression abolishes cycle-dependent changes.

---

Neuroscientists have plumbed the depths of the mind and brain to extraordinary lengths, but occasionally we forget that the brain is part of a larger, integrated biological system. The brain is an endocrine organ whose structure and function is intimately tied to the action of neuromodulatory hormones (1–4). Sex steroid hormone receptors are expressed throughout the brain, with high expression in the hippocampus and surrounding medial temporal lobe (MTL) (5–7). Rodent (1, 2, 8, 9) and non-human primate (10) studies have established 17β-estradiol and progesterone as powerful regulators of hippocampal subfield morphology. For example, dendritic spine density in CA1 pyramidal neurons fluctuates by 30% across the rat estrous cycle (1, 11). Estradiol enhances synaptic plasticity and spine proliferation in rodent CA1 neurons (3, 10, 12, 13) and progesterone inhibits this effect (1). Less is known about the influence of sex hormones on human MTL subregions. The extent of our understanding comes from observations of gross hippocampal volume differences in discrete stages of the menstrual cycle (14, 15) and emerging evidence that hippocampal volume declines following spontaneous (16) and surgical (17) menopause.

A central feature of the mammalian endocrine system is that hormone secretion varies over time. Circadian, infradian, and circannual rhythms are essential for sustaining many physiological processes. During an average human menstrual cycle, spanning 25–32 days, women experience a ∼12-fold increase in estradiol and an ∼800-fold increase in progesterone. Despite this striking change in endocrine status, we lack a complete understanding of how MTL structures respond to rhythmic changes in sex hormone production over time. The study of brain–hormone interactions in human neuroscience relies heavily on cross-sectional designs that, by nature, cannot capture dynamic changes in hormone production. However, an emerging trend in human neuroimaging is to flip the cross-sectional model by densely sampling individuals over timescales of days, weeks, months, or years to provide greater insight into the dynamic properties of the human brain (18–20).

Although most human MRI studies treat the hippocampus as a homogeneous structure, it is comprised of distinct subfields (cornu ammonis 1-4, dentate gyrus, and subiculum) each with a unique cytoarchitecture and circuitry. The hippocampus proper is surrounded by highly interconnected structures (entorhinal, perirhinal, and parahippocampal cortex) that, together, comprise the medial temporal lobe (MTL). Advances in high resolution MRI allow for the *in vivo* measurement of hippocampal subfield volume (21). MTL subregions are differentially impacted by chronological aging (22, 23) and harbor distinct patterns of neuropathology in the progression to Alzheimer’s disease (24) However, very little is known about sex hormones’ impact on unique MTL subregions, since the preclinical animal literature is focused predominantly on CA1 (with proliferative effects of estradiol and suppressive effects of progesterone on synaptic plasticity) while human studies are limited to estimates of total hippocampal volume (with larger volumes generally coinciding with periods of enhanced estrogenicity (14, 15). This represents a major gap in our understanding of the spatial and temporal pattern with which sex hormones shape MTL morphology.

In a series of dense-sampling studies, we used high-resolution imaging of the MTL to determine whether variation in sex hormone concentrations over time impacts MTL morphology with subregion resolution. First, we assessed the relationship between MTL subregion volume and endogenous fluctuations in sex hormones across a complete menstrual cycle (Study 1, n = 30 days). Next, we determined the effects of chronically and selectively suppressing progesterone over the same timescale (Study 2, n = 30 days). This longitudinal approach affords direct comparisons between freely-cycling and hormonally-suppressed states within the same individual, providing powerful insight into the relationship between sex hormones and MTL morphology.

## Methods

### Participant

A healthy, female (author L.P., age 23) participated in two longitudinal, dense-sampling, deep-phenotyping studies. The participant had no history of neuropsychiatric diagnosis, endocrine disorders, or prior head trauma and no history of smoking. She had a history of regular menstrual cycles (no missed periods, cycle occurring every 26–28 days) and had abstained from hormone-based medications for 12 months preceding Study 1. The participant gave written informed consent and the study was approved by the University of California, Santa Barbara Human Subjects Committee.

### Study design

To map MTL subregion volume in the human brain across a menstrual cycle, the participant underwent daily testing for 30 consecutive days while freely cycling (Study 1), with the first test session determined independently of cycle stage for maximal blindness to hormone status. In a follow-up study (Study 2) which took place 12 months after Study 1, the participant repeated the 30-day protocol while on a hormone regimen (0.02 mg ethinyl-estradiol, 0.1 mg levonorgestrel, Aubra, Afaxys Pharmaceuticals), which she began 10 months prior to the start of data collection. Procedures for Studies 1 and 2 were identical (**Fig. S1**): the participant underwent a time-locked daily blood draw immediately followed by an MRI scan. Endocrine samples were collected, at minimum, after two hours of no food or drink consumption (excluding water). The participant refrained from consuming caffeinated beverages before each test session. The MRI session lasted one hour and consisted of structural and functional MRI sequences (functional data are reported elsewhere). Neuroimaging data will be publicly accessible upon publication.

### Endocrine procedures

A licensed phlebotomist inserted a saline-lock intravenous line into the hand or forearm daily to evaluate hypothalamic-pituitary-gonadal axis hormones, including serum levels of gonadal hormones (17β-estradiol and progesterone) and the pituitary gonadotropins luteinizing hormone (LH) and follicle stimulating hormone (FSH). One 10 ml blood sample was collected in a vacutainer SST (BD Diagnostic Systems) each session. The sample clotted at room temperature for 45 minutes until centrifugation (2000 × g for 10 minutes) and were then aliquoted into three 1 ml microtubes. Serum samples were stored at -20°C until assayed. Sex steroid concentrations were determined via liquid chromatography-mass spectrometry (LC-MS) at the Brigham and Women’s Hospital Research Assay Core (BRAC). Assay sensitivities, dynamic range, and intra-assay coefficients of variation were as follows: *estradiol*: 1.0 pg/ml, 1–500 pg/ml, <5% relative standard deviation (RSD); *progesterone*: 0.05 ng/ml, 0.05–10 ng/ml, 9.33% RSD; *testosterone*: 1.0 ng/dL, 1–2000 ng/dL, <4% RSD. The gonadotropins FSH and LH levels were determined via chemiluminescent assay (Beckman Coulter) at the BRAC. The assay sensitivity, dynamic range, and the intra-assay coefficient of variation were as follows: *FSH*: 0.2 mIU/ml, 0.2–200 mIU/ml, 3.1–4.3%; *LH*: 0.2 mIU/ml, 0.2–250 mIU/ml, 4.3–6.4%.

### MRI Acquisition

The participant underwent a daily magnetic resonance imaging (MRI) scan on a Siemens 3T Prisma scanner equipped with a 64-channel phased-array head/neck coil (of which 50 coils are used for axial brain imaging). High-resolution anatomical scans were acquired using a *T*_*1*_-weighted magnetization prepared rapid gradient echo (MPRAGE) sequence (TR = 2500 ms, TE = 2.31 ms, T_1_ = 934 ms, flip angle = 7°, 0.8 mm thickness) followed by a gradient echo fieldmap (TR = 758 ms; TE_1_ = 4.92 ms; TE_2_ = 7.38 ms; flip angle = 60°). A *T*_2_-weighted turbo spin echo (TSE) scan was also acquired with an oblique coronal orientation positioned orthogonally to the main axis of the hippocampus (TR/TE= 8100/50 ms, flip angle = 122°, 0.4 × 0.4 mm^2^ in-plane resolution, 2 mm slice thickness, 31 interleaved slices with no gap, total acquisition time = 4:21 min). To minimize motion, the head was secured with a custom, 3D-printed foam head case (https://caseforge.co/) (days 8–30 of Study 1 and days 1–30 of Study 2), sandbag cushions were placed on the participant’s legs, and a strap was placed over the shoulders. Overall motion (mean framewise displacement) on a preceding functional scan was negligible, with less than 150 microns of motion on average each day (**Fig. S2**).

### Hippocampal Segmentation

*T*_1_- and *T*_2_-weighted images (n = 60) were submitted to the automatic segmentation of hippocampal subfields package (ASHS) (21) for bilateral parcellation of seven MTL subregions: CA1, CA2/3, dentate gyrus (DG), subiculum (SUB), perirhinal cortex (PRC), entorhinal cortex (ERC), and parahippocampal cortex (PHC) (**Fig. 1B**). The ASHS segmentation pipeline automatically segmented the hippocampus in the *T*_2_-weighted MRI scans using a segmented population atlas, the Princeton Young Adult 3T ASHS Atlas template (n = 24, mean age 22.5 years (25)). A rigid-body transformation aligned each *T*_2_ image to the respective *T*_*1*_ scan for each day. Using Advanced Normalization Tools (ANTs) deformable registration, the *T*_*1*_ was registered to the population atlas. The resulting deformation fields were used to resample the data into the space of the left and right template MTL regions of interest (ROI). Within each template ROI, each of the *T*_2_-weighted scans of the atlas package was registered to that day’s *T*_2_ scan. The manual atlas segmentations were then mapped into the space of the *T*_2_ scan, with segmentation of the *T*_2_ scan computed using joint label fusion. Finally, the corrective learning classifiers contained in the atlas package were applied to the consensus segmentation produced by joint label fusion. The output of this step is a corrected segmentation of the *T*_2_-weighted scan. Further description of the ASHS protocol can be found in (21).

**Figure 1.**
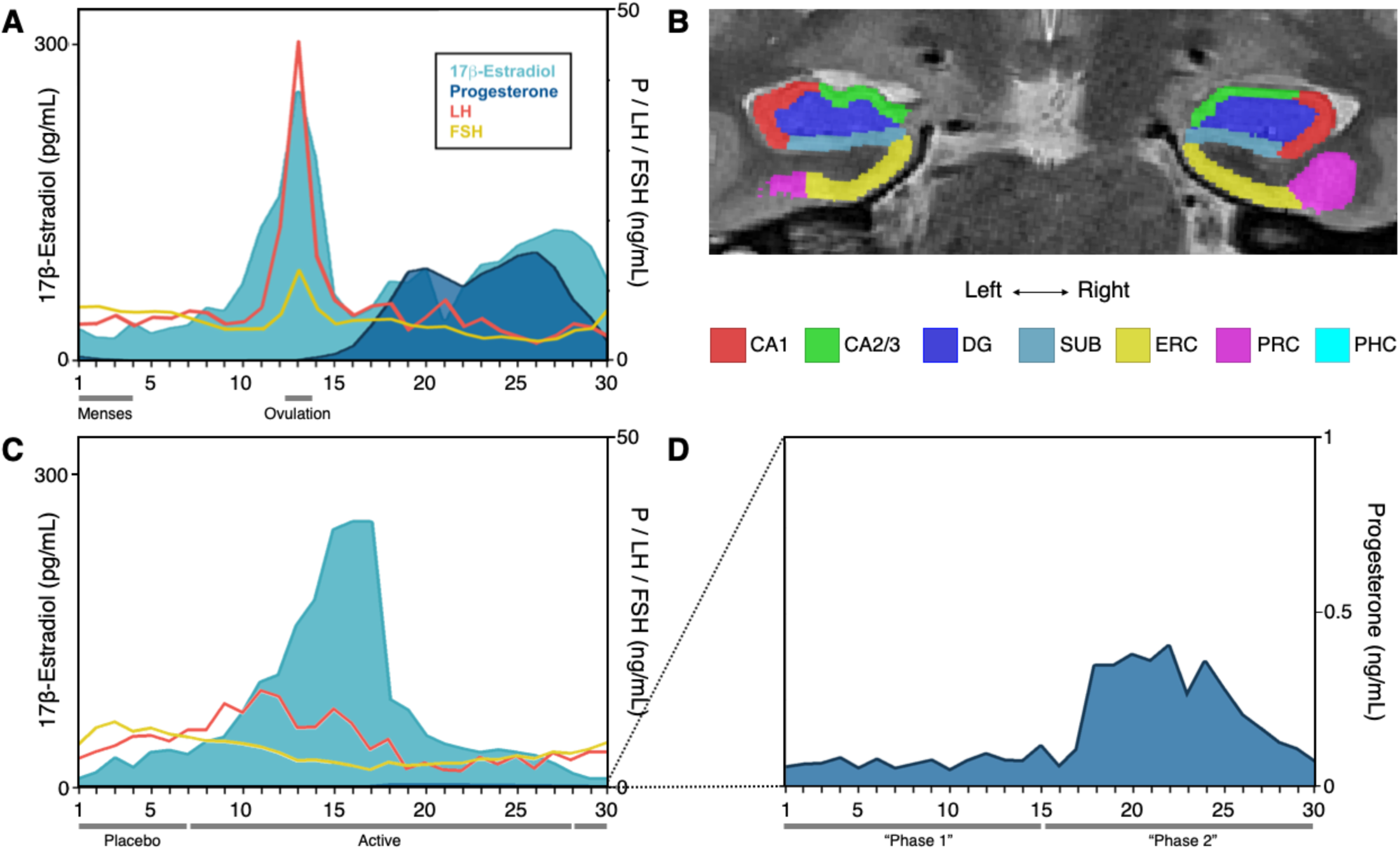
Pituitary and gonadal hormones by experiment and subregion parcellations. **A**. Pituitary gonadotropins (LH, FSH) and gonadal hormones (estradiol, progesterone) across 30 days of a complete menstrual cycle. Estradiol exhibits a 12-fold increase prior to ovulation. Progesterone concentrations increase 800-fold during the luteal phase of the natural cycle. **B**. Sample slice with Automatic Segmentation of Hippocampal Subfields (ASHS)-segmented hippocampal subfields and MTL cortex labels. **C**. Hormone concentrations across the 30-day hormone suppression study. Estradiol concentrations were unmodified, while progesterone was suppressed by 97% on average. **D**. Progesterone concentrations during the hormone suppression study plotted on a 1 ng/ml scale to illustrate some preserved modulation of progesterone, with a 10-fold rise across days 16–23. Note that ‘Day 1’ indicates first day of menstruation, not first day of experiment. Experiment day 1 corresponded to cycle day 21 in Study 1 and cycle day 18 in Study 2. Abbreviations: P, progesterone; *LH*, luteinizing hormone; FSH, follicle-stimulating hormone; CA, cornu ammonis; DG, dentate gyrus; SUB, subiculum; ERC, entorhinal cortex; PRC, perirhinal cortex; PHC, parahippocampal cortex.

*T*_*2*_ scans and segmentations were first visually examined using ITK-SNAP (26) for quality assurance and then subjected to more fine-grained manual editing in native space using ITK-SNAP (v.3.8.0-*b*). One bilateral segmentation (Study 2, Day 26) was removed as an outlier due to significant underestimation of volumes across subregions (>3x the standard deviation from the mean). Anterior and posterior hippocampal ROIs, which corresponded to the anterior head and tail, were integrated into the label output in keeping with the Olsen-Amaral-Palombo (OAP) segmentation protocol (27) to ensure full coverage of the hippocampus when computing total hippocampal volume. The anterior extent of the segmented labels was anchored 4 mm (2 slices) anterior to the appearance of the limen insulae, and the posterior extent was anchored to the disappearance of hippocampal gray matter from the trigone of the lateral ventricle. The boundary between anterior and posterior parahippocampal gyrus was determined by ASHS and not edited. In instances where automatic segmentation did not clearly correspond to the underlying neuroanatomy, such as when a certain label was missing a number of gray matter voxels, manual retouching allowed for individual voxels to be added or removed (**Fig. S3)**. All results are reported using the manually retouched subregion volumes to ensure the most faithful representation of the underlying neuroanatomy. Note that the segmenter (C.T.) was blind to the hormone status of the participant. Scans were randomized and segmentation was performed in a random order, not by day of experiment.

Regarding reliability, the ASHS pipeline has reported a cross-validation of 0.779 between automatic and manual segmentations (21), though we acknowledge that this may differ between atlases and datasets, and the Princeton atlas utilized for these analyses does not report cross-validation values. To assess intra-rater reliability for the present analyses, five days were randomly selected to undergo manual editing a second time. The generalized Dice similarity coefficient (28) across subregions was 0.98 and the Intraclass Correlation Coefficient was 0.998, suggesting robust reliability in segmentation.

### Gray Matter Volume

#### Whole-brain

Total whole-brain gray matter volume (wbGMV) for each study session was calculated using the Segment Longitudinal Data module in the cat12 toolbox in SPM12 (29). Statistical models evaluating the relationship between MTL subregion volume and endocrine values are reported with and without adjustment for wbGMV, which had little bearing on the overall results.

#### Precentral gyrus

Bilateral precentral gyrus (PreCG) volumes were specified using the automated anatomical labelling (AAL) atlas (30) and extracted using the MarsBaR (31) toolbox in SPM12. Bilateral PreCG was identified as a control region of interest to evaluate the relationship between progesterone and a brain region not known to be strongly influenced by sex steroid hormones.

### Statistical Analysis

All analyses were conducted in *R* (version 3.4.4). To evaluate the relationship between subregion volumes and endogenous sex hormones across the menstrual cycle, linear regression was conducted between bilateral subregion volumes (average of left and right) and concentrations of both 17β-estradiol and progesterone. We had no strong *a priori* hypothesis that structure– hormone relationships would differ by hemisphere; thus, volumes are reported averaged across hemispheres. For completeness, values by hemisphere are provided in **Supplementary Results (Table S2)**. Note that results by hemisphere do not differ substantially from findings reported below. Relationships were considered significant only if they met the Bonferroni correction for multiple comparisons of *p* < .007 (*p* < .05/7 regions). Though volumes were normally distributed (Shapiro–Wilk *p* > .07) across subregions, progesterone values were not, so these relationships were further investigated using non-parametric Spearman rank correlation. Results were highly consistent between the two approaches.

Next, wbGMV was added to the hormone–volume linear models to determine whether the relationship between progesterone and subregion volume held after accounting for variance attributable to wbGMV. Similarly, the number of slices attributed to a subregion can vary by scan. Given the shared boundary between anterior (PHC, ERC) and posterior (PHC) subregion volumes, shifts in slice number between these regions (e.g. an additional slice attributed to PHC might correspond with one fewer slice attributed to PRC and ERC) could contribute to the inverse relationships observed between progesterone and anterior and posterior subregion volumes. To determine the extent to which these inverse relationships were attributable to differences in slice number, statistical models evaluating the relationship between MTL subregion volume and endocrine values are provided with adjustment for slice number. Note that results did not differ appreciably across models.

MTL subregions are highly inter-connected. To help clarify the specificity of progesterone’s effects, we examined a region of interest outside the MTL (one not thought to be affected by fluctuations in sex hormones), the PreCG. As before, we performed linear regressions between bilateral PreCG volume and progesterone.

Progesterone displayed a bimodal distribution with low concentrations in the follicular phase and high concentrations in the luteal phase (Study 1, **Fig. 1A**), with an 800-fold change between lowest to highest values. To further characterize the data, we split the 30-day natural cycle into two phases corresponding to the first and second half of the study period: “Phase 1” (cycle days 1–15, *M*_P1_ 0.212 ng/ml, *range* 0.019–0.97 ng/ml) and “Phase 2” (cycle days 16–30, *M*_P2_ 10.074 ng/ml, *range* 2.13–15.5 ng/ml). Two-sample *t*-tests were conducted to compare subregion volumes during Phases 1 and 2.

In the follow-up study (Study 2; preregistered under (32), the participant was placed on a hormonal regimen that selectively suppressed circulating progesterone (**Fig 1C,D**). We tested the hypothesis that holding progesterone concentrations chronically low (comparable to levels observed in the early follicular phase of the natural cycle) would abolish the progesterone-mediated effects observed in Study 1. Two-factor ANOVAs were conducted to compare the main effects of hormone status (naturally cycling vs. hormonally suppressed), progesterone phase (Phase 1 vs. Phase 2) and their interaction on MTL subregion volume. Post-hoc comparisons were conducted using Tukey’s HSD tests.

## Results

### Endocrine assessments

Analysis of daily sex hormone (by LC-MS) and gonadotropin (by chemiluminescent immunoassay) concentrations across the menstrual cycle (Study 1) confirmed the expected rhythmic changes of a typical cycle (33), with a total cycle length of 27 days. Serum levels of estradiol were lowest during menses (days 1–4) and peaked in the late follicular phase (day 13). Progesterone concentrations showed the expected dramatic rise during the mid-luteal phase (**Fig. 1A; Table S1)** and surpassed 5 ng/ml, signaling an ovulatory cycle (34).

In Study 2, the participant was placed on a daily low-dose regimen of 0.02 mg ethinyl-estradiol, 0.1 mg levonorgestrel, which selectively suppressed circulating progesterone. The concentration and dynamic range of 17β-estradiol was similar during progesterone suppression (*M* = 66.2 pg/ml, *range*: 5–246 pg/ml) compared to naturally cycling (*M =* 82.8 pg/ml, *range* 22–264 pg/ml; *t*(57) = 0.18, *p* = .24). In contrast, progesterone concentrations were suppressed by ∼97% on average (Study 1: *M* = 5.14 ng/ml, *range* = 5–15.5 ng/ml; Study 2: *M* = 0.15 ng/ml, *range* = 0.04–0.40; *t*(57) = 4.58, *p* < .0001) (**Fig. 1C; Table S1**). Note that LC-MS assessments of exogenous hormone concentrations (attributable to the hormone regimen itself) showed that serum concentrations of ethinyl estradiol were very low (*M* = 0.01 ng/ml; *range* 0.001–0.016 ng/ml) and below 1.5 ng/ml for levonorgestrel (*M* = 0.91 ng/ml; *range* = 0.03–1.43 ng/ml).

### Intrinsic fluctuations in sex hormones and MTL subregion volume across a menstrual cycle

To begin, we tested the hypothesis that MTL subregion volume is associated with intrinsic fluctuations in estradiol and progesterone across a complete menstrual cycle (n = 30 days). Results reported below met a Bonferroni corrected threshold of significance of *p* = .007. We observed no significant relationship between 17β-estradiol and volume (all *p* > .14). Progesterone, by contrast, showed significant associations with subregions throughout the MTL. Progesterone was positively associated with GMV in CA2/3 (F_(1,28)_15.37, p < .001, R^2^_adjusted_ = .33) and parahippocampal cortex (F_(1,28)_19.06, p < .001, R^2^_adj_ = .38) (**Fig. 2A, Table 1**); it was negatively associated with GMV in perirhinal cortex (F_(1,28)_16.33, p < .001, R^2^_adj_ = .35) and entorhinal cortex (F_(1,28)_18.82, p < .001, R^2^_adj_ = .38) (**Fig. 3A, Table 1, Supplementary Fig. 5**). This pattern of results was robust and did not change when computing non-parametric Spearman rank-coefficient correlations (all *p* ≤ .005). Next, multiple regression confirmed that these relationships remained significant after accounting for wbGMV (CA2/3: *β* = .54, *p* = .003; PHC: *β* = .64, *p* < .001; PRC: *β* = -.61, *p* < .001; ERC: *β* = -.60, *p* < .001); and after accounting for slice number (CA2/3: *β* = .64, *p* < .001; PHC: *β* = .51, *p* < .001; ERC: *β* = -.33, *p* = .02) with the exception of PRC volume (*β* = -.03) which was no longer significantly predicted by progesterone. Results did not differ when both wbGMV and slice number were included in the same model. Dividing the menstrual cycle into Phase 1 (days 1–15, low progesterone) and Phase 2 (days 16–30, high progesterone) further confirmed differences in subregion volume as a function of progesterone (CA2/3: *t*(28) = -3.85, *p* < 0.001; PHC: *t*(28) = -2.95, *p* < 0.01; PRC: *t*(28) = 2.90, *p* < 0.01; ERC: *t*(28) = 2.34, *p* < 0.03) (**Fig. 2B, 3B**). We did not observe a relationship between whole hippocampal volume (sum of subfield as well as anterior and posterior volumes) and estradiol or progesterone, nor did we observe a significant relationship between bilateral PreCG volume and progesterone (*p* = .44).

**Table 1.**
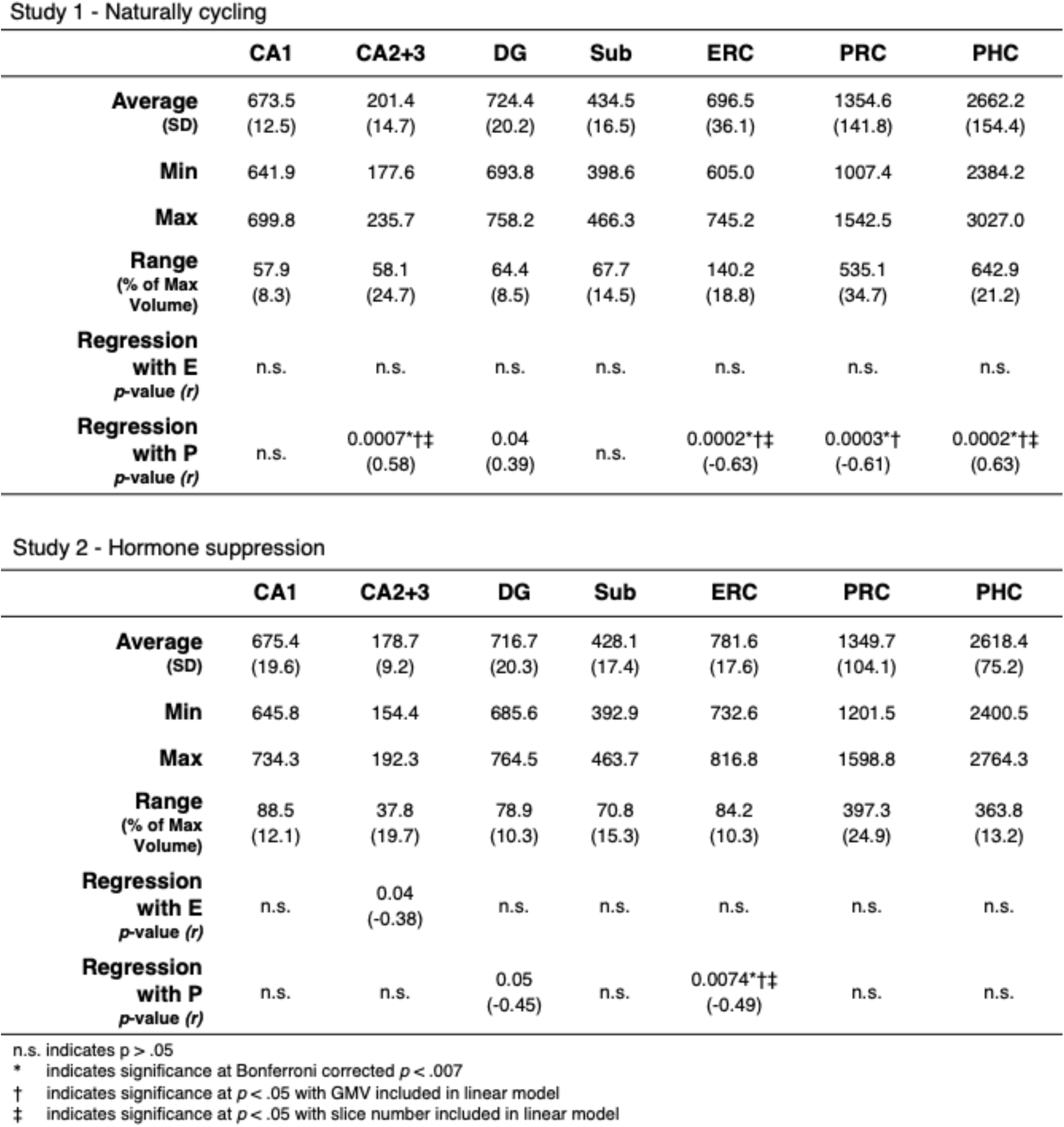
Medial temporal lobe volumes (mm^3^) and association with sex hormones by experiment

**Figure 2.**
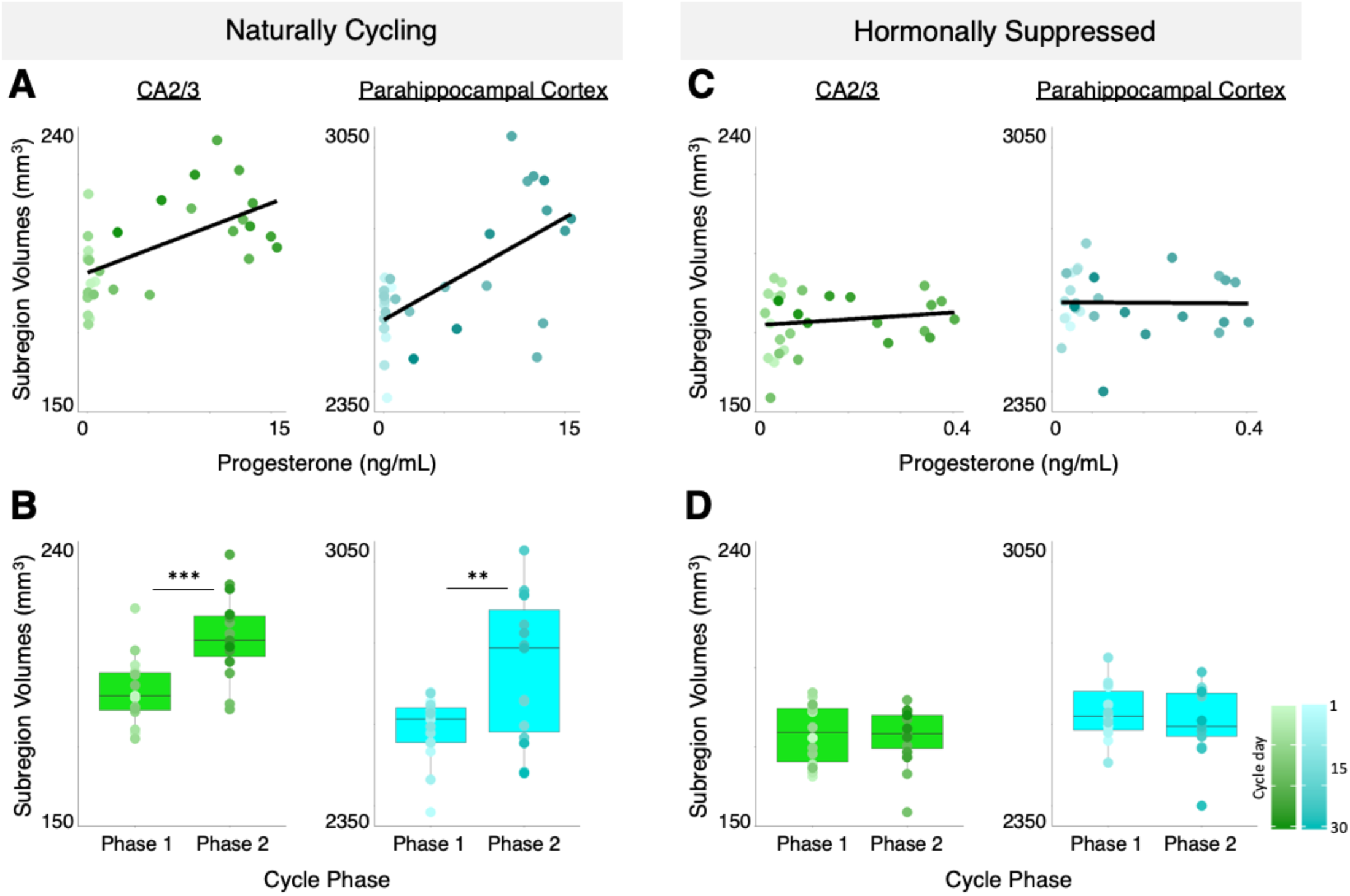
Relationship between progesterone concentrations and MTL subregion volume across the menstrual cycle and during hormone suppression. **A**. In CA2/3 and parahippocampal cortex, GMV was *positively* correlated with progesterone and **B**. significantly different between Phase 1 (low progesterone) and Phase 2 (high progesterone) across the cycle. **C-D**. This modulation was abolished during chronic hormone suppression (Study 2). Comparisons between the two studies (**B, D**) reveal that GMV in CA2/3 was reduced following hormone suppression relative to freely cycling (*p* <.0001; effect size, η^2^ =.47). Individual data points are color-coded to indicate cycle day, from day 1 (light circles) to day 30 (dark circles). \*\**p* < .01, \*\*\**p* < .001; GMV, gray matter volume.

**Figure 3.**
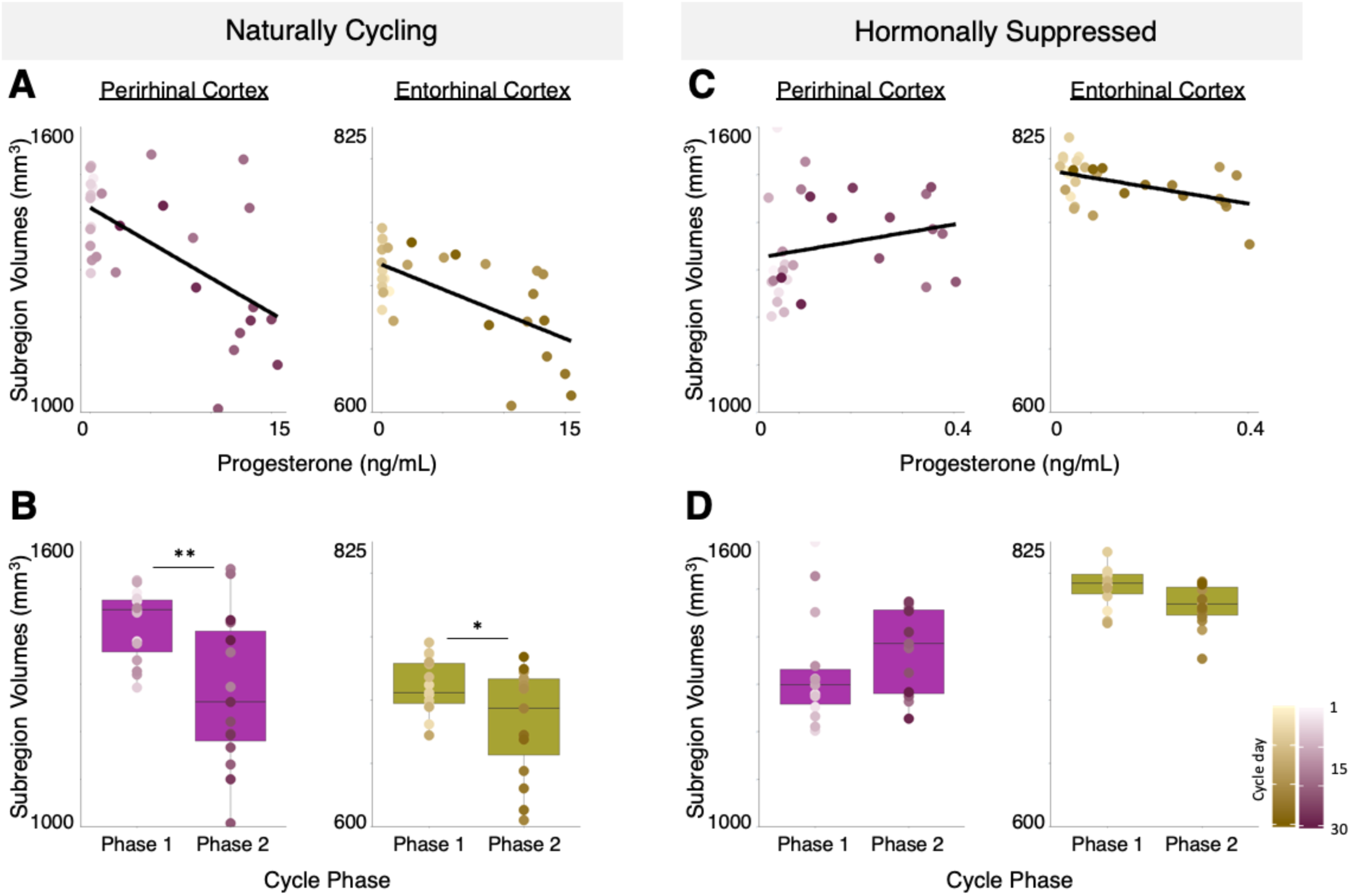
Relationship between progesterone concentrations and MTL subregion volume across the menstrual cycle and during hormone suppression **A**. In perirhinal and entorhinal cortex, GMV was *negatively* associated with progesterone and **B**. significantly different between Phase 1 (low progesterone) and Phase 2 (high progesterone) across the cycle. This modulation was not observed under chronic hormone suppression (**C, D**). Comparisons across the two studies (**B, D**) reveal that GMV in entorhinal cortex was larger during hormone suppression relative to freely cycling (*p* <.0001; effect size, η^2^ =.70). Individual data points are color-coded to indicate cycle day, from day 1 (light circles) to day 30 (dark circles). *p < .05, **p < .01; GMV, gray matter volume.

### Impact of hormone suppression on MTL subregion volume

Next, to determine whether the changes in MTL subregion volumes observed in Study 1 were dependent on dynamic fluctuations in progesterone, we repeated the experiment with progesterone concentrations chronically and selectively suppressed. Estradiol concentrations varied as before (**Fig. 1C**); however, progesterone concentrations were suppressed by ∼97% on average. Under hormone suppression, progesterone concentrations were comparable to values observed in the early follicular phase of the natural cycle, effectively reducing the 800-fold rise in progesterone that typically occurs in the luteal phase (**Fig. 1A**) to a 10-fold change (**Fig. 1C,D**). The 30-day experimental period was divided into “Phase 1” and “Phase 2”, as before, to capture the small degree of progesterone modulation that remained (**Fig. 1D**). As above, significance was held to a Bonferroni corrected threshold of *p* < .007.

When progesterone concentrations were chronically suppressed, the previously observed relationships between progesterone and subregion volumes were abolished or markedly reduced (**Fig. 2C,D; Fig. 3C,D**). Regions that showed strong progesterone-dependent volumetric changes during the menstrual cycle (CA2/3, PHC and PRC) displayed no such relationship under progesterone suppression (all *p* > 0.25). The single exception was area ERC, which continued to show a negative relationship with progesterone (F_(1,27)_8.39, *p* = .007, R^2^_adj_ = .21).

Two-way ANOVAs were conducted to directly compare subregion volumes across studies as a function of progesterone phase (low, high) and hormone condition (naturally cycling, hormonally suppressed). In CA2/3 there were significant main effects of progesterone phase (F_(1,55)_7.47; *p* = .008; effect size, η^2^ =.054) and hormone condition (F_(1,55)_65.0; *p* < .0001; effect size, η^2^ =.473), as well as a significant interaction (F_(1,55)_9.93; *p* = .003; effect size, η^2^ =.072).

Post-hoc tests established that CA2/3 volumes were significantly different between high progesterone (*M* = 210.1 mm^3^) and low progesterone (*M* = 193.1 mm^3^) conditions in Study 1 (*p* < .001), but not in Study 2 (*M* _high P_ = 180.0 mm^3^, *M* _low P_ = 177.6 mm^3^; *p* = .99). Finally, CA2/3 volumes were smaller overall under hormone suppression conditions (*M =* 178.7 mm^3^) relative to naturally cycling conditions (*M =* 201.4 mm^3^, *p* < .0001) (**Table 2**).

**Table 2.** Comparison of subfield volumes – Naturally cycling vs. Hormonally suppressed

|  | CA1 | CA2+3 | DG | Sub | ERC | PRC | PHC |
| --- | --- | --- | --- | --- | --- | --- | --- |
| **Naturally Cycling**<br>**Average**<br>**(SD)** | 673.5<br>(12.5) | 201.4<br>(14.7) | 724.4<br>(20.2) | 434.5<br>(16.5) | 696.5<br>(36.1) | 1354.6<br>(141.8) | 2662.2<br>(154.4) |
| **Hormonally Suppressed**<br>**Average**<br>**(SD)** | 675.4<br>(19.6) | 178.7<br>(9.2) | 716.7<br>(20.3) | 428.1<br>(17.4) | 781.6<br>(17.6) | 1349.7<br>(104.1) | 2618.4<br>(75.2) |
| **Effect size, $\eta^2$** |  | .47 |  |  | .70 |  |  |
| **p-value** | n.s. | < 0.0001 | n.s. | n.s. | < 0.0001 | n.s. | n.s. |

n.s. indicates $p > 0.05$

In PHC, we observed a significant main effect of progesterone level, (F_(1,55)_4.92; *p* = .03; effect size, η^2^ = .070) and an interaction between progesterone level and hormone condition (F_(1,55)_8.46; *p* = .005 effect size, η^2^ = .119). The main effect of hormone condition was not significant, indicating no overall difference in PHC volume between Studies 1 and 2. Post-hoc tests established that PHC volumes were impacted by variation in progesterone in Study 1 (*M* _high P_ = 2736.2 mm^3^, *M* _low P_ = 2588.2 mm^3^, *p* < .005)— an effect that was abolished in Study 2 (*M* _high P_ = 2602.5 mm^3^, *M* _low P_ = 2633.2 mm^3^, *p* = .96).

In PRC, only the interaction between progesterone level and hormone status was significant (F_(1,55)_9.36; *p* = .003; effect size, η^2^ =.139). Post-hoc tests again showed that this interaction was driven by progesterone exerting its effect on PRC volume in Study 1 (*M* _high P_ = 1280.7 mm^3^, *M* _low P_ = 1414.0 mm^3^, *p* < .01) but not in Study 2 (*M* _high P_ = 1377.2 mm^3^, *M* _low P_ = 1324.0 mm^3^, *p* =.74).

Finally, in ERC, main effects of progesterone level (F_(1,55)_8.60; *p* = .005; effect size, η^2^ =.040) and hormone status (F_(1,55)_152.26; *p* < .0001; effect size, η^2^ =.702) were significant, with no interaction. There was a significant inhibitory effect of progesterone on ERC volumes in Study 1 (*M* _high P_ = 682.1 mm^3^, *M* _low P_ = 710.9 mm^3^, *p* = .023), but not Study 2 (*M* _high P_ = 774.4 mm^3^, *M* _low P_ = 788.2 mm^3^, *p* = .46), though the direction of the effects were similar. ERC volumes were significantly larger under hormone suppression conditions (*M* = 781.6 mm^3^) compared to naturally cycling (*M* = 696.5 mm^3^, *p* < .0001) (**Table 2**).

## Discussion

In a series of dense-sampling, deep-phenotyping studies, we tested the hypothesis that intrinsic fluctuations in sex hormones impact MTL subregion volume across the human menstrual cycle, and that chronic hormone suppression abolishes this effect. In Study 1, a naturally-cycling female underwent high resolution imaging of the MTL and venipuncture for 30 consecutive days, capturing the dynamic endocrine changes that unfold over a complete menstrual cycle. Endogenous fluctuations in progesterone impacted gray matter volume in several subregions of the medial temporal lobe, including CA2/3, parahippocampal, entorhinal and perirhinal cortex. In Study 2, the experiment was repeated while the participant was on a hormone regimen that chronically and selectively suppressed circulating progesterone. The progesterone-mediated effects observed in Study 1 were no longer evident, and volumetric changes in CA2/3 and entorhinal cortex were observed during chronic progesterone suppression relative to freely cycling conditions. Together, these results reveal progesterone’s ability to dynamically shape medial temporal lobe morphology across the menstrual cycle.

### Sex hormones influence hippocampal morphology across species

Three decades of rodent and nonhuman primate studies offer unambiguous evidence that sex steroid hormones influence the synaptic organization of the hippocampus (1, 3, 35–38). At the epigenetic level, sex hormones induce chromatin modifications that promote hippocampal plasticity (39). At the synaptic level, sex hormones regulate spine proliferation in the hippocampus (3). In a series of seminal findings, Woolley and colleagues (11, 12) discovered that dendritic spine density in CA1 neurons varies by ∼30% over the 4–5-day rodent estrous cycle. Similarly, hormone deprivation (via gonadectomy) in the rat (1, 40) and African green monkey (41) leads to a pronounced loss of spines on CA1 neurons, which is reversed by hormone replacement. Finally, at the macroscopic level, total hippocampal volume is regulated by sex hormones in the meadow vole (42) and fluctuates across the estrous cycle in the mouse (43). *In-vivo* MRI in mice demonstrates that hormone-mediated hippocampal changes can occur rapidly, with differences detectable within a 24-hour period (43). Together, these findings provide converging evidence that sex hormones can induce structural changes in the hippocampus on a rapid timescale.

Human studies offer further evidence that ovarian status impacts hippocampal morphology. The abrupt hormonal changes associated with early surgical menopause lead to structural changes in the medial temporal lobe, including thinning of the parahippocampus/entorhinal cortex (17), while hormone supplementation in postmenopausal women increases hippocampal volume (44). In pregnancy, the rise in sex hormone production across the gestational period impacts hippocampal plasticity in rodents (45–47) and likely mediates the decline in hippocampal volume that occurs during pregnancy in humans (48). Our data extend this literature by establishing volumetric changes in the medial temporal lobe across the ∼28-day menstrual cycle.

### Sex hormones regulate hippocampal morphology across the human menstrual cycle

A handful of cross-sectional and sparse-sampling studies have probed hippocampal volume at discrete stages of the menstrual cycle (14, 15, 49). Protopopescu et al. (14) reported greater right anterior hippocampal volume in women during the follicular versus luteal phase. Similarly, Pletzer et al. (49) observed greater gray matter volume in the parahippocampal gyrus during the early follicular versus mid-luteal phase. One study of naturally cycling women captured four discrete stages of the menstrual cycle, reporting a positive correlation between estrogen and parahippocampal gyrus volume (15). By distilling the menstrual cycle into a few timepoints, these approaches, though valuable, may not capture the full impact of sex hormones over time. Cross-sectional and sparse-sampling approaches obscure the rhythmic production of sex hormones over a complete cycle. A formative study by Barth and colleagues (50) densely sampled a woman every few days across the menstrual cycle, observing a relationship between estradiol and gray matter changes in the left hippocampus using voxel-based morphometry estimates. Here, we used high resolution MTL imaging to show that medial temporal lobe structures respond uniquely to dynamic changes in sex hormone production.

Based on the existing literature, our *a priori* hypotheses anticipated a positive relationship between estradiol and CA1 volume across the menstrual cycle; however, this was not borne out in the data. No relationship with estradiol was significant. We also predicted a suppressive effect of progesterone on volume, which was observed in ERC and PRC. More surprisingly, progesterone fluctuations across the cycle were positively related to CA2/3 and PHC volume. Note that no relationship was observed between progesterone and precentral gyrus volume, a control region outside of the MTL not expected to show strong hormone-dependent effects. To our knowledge, this is the first longitudinal study of subfield volume changes across the menstrual cycle and in response to chronic progesterone suppression. Future studies should build on these findings using high-field imaging to interrogate subfield volume changes triggered by hormone suppression and subsequent hormone add-back protocols. Doing so will help clarify mixed findings and deepen our understanding of sex hormone action at the subfield level.

### Potential neurobiological mechanisms of progesterone’s influence in MTL

Woolley and McEwen (1) discovered that progesterone has a biphasic effect on spine density in the hippocampus, eliciting an initial increase (within 2–6 hours) followed by a sharp and sustained decrease. Progesterone administration triggers the regression of dendritic spines in rodent CA1 neurons, an effect that is blocked by progesterone receptor antagonists. The changes we observed in MTL volume in the human brain may partially reflect hormone-driven changes in dendritic spines or synapse density. However, given the difference in scale, it seems unlikely that changes at the synaptic level could fully account for volumetric changes at the subregion level, and may be why we did not observe changes in CA1. To our knowledge, no animal study has directly linked observations made at the mesoscopic scale of regional volume to the microscopic scale of synapse/spine morphology. Further, sex hormones’ ability to regulate hippocampal structural plasticity has been carefully studied in rodent CA1 and dentate gyrus, while much less is known about other MTL regions. Future rodent and nonhuman primate studies are warranted to resolve these outstanding questions.

In the present study, progesterone’s impact on MTL volume was subregion-specific, with proliferative effects in CA2/3 and parahippocampal cortex and regressive effects on entorhinal and perirhinal cortex. Evidence for subregion-specific effects of progesterone comes from the study of neurotrophins. Progesterone blocks estrogen’s ability to upregulate growth factors (including nerve growth factor, brain derived neurotrophic factor and neurotrophin 3) in entorhinal cortex but not in hippocampal subfields (35). In our study, increased progesterone across the cycle was associated with decreased ERC volume. When progesterone levels were chronically blunted in Study 2, ERC volume rebounded, perhaps because progesterone’s inhibitory action in this region had been expunged. Subregion-specific morphological changes could also stem from progesterone’s ability to modulate neural progenitor cell proliferation. For example, progesterone blocks estradiol’s enhancement of cell proliferation in the dentate gyrus (51). Thus, progesterone’s effects within the MTL may be mediated by the receptor expression profile and proliferative capacities of different subregions. Differences in receptor expression could produce a heightened sensitivity to progesterone (or its metabolites) in unique MTL regions. Although the presence of classical nuclear progesterone receptors in the mammalian hippocampus is well-established (52), *in vivo* animal studies and *ex-vivo* human studies are needed to determine distinct receptor expression profiles at the subregion level and to quantify the relative concentration of progesterone and progesterone metabolites throughout the MTL. Finally, *de novo* production of neurosteroids in the hippocampus are important drivers of synaptic plasticity. Although speculative, a sustained decrease in ovarian-derived steroid hormones could trigger a compensatory upregulation of brain-derived neurosteroids in a subfield-specific manner, contributing to blood hormone/brain volume relationships that are unique across MTL regions.

In addition to its neuronal effects, progesterone also impacts non-neuronal cell populations. Progesterone increases the number of oligodendrocytes expressing myelin-specific proteins (53) and promotes myelination in Schwann cells (54) on the order of days (55) and weeks (54). While this has been observed during development and in response to injury (reviewed in (56, 57), the effects of progesterone on myelination across the menstrual cycle are not yet known. In a diffusion tensor imaging study, Barth and colleagues (50) observed changes in fractional anisotropy (a purported measure of white matter structural integrity) in the hippocampus across the menstrual cycle, although no associations with progesterone were found.

The progesterone-dependent effects we observed could also be mediated indirectly by progesterone metabolites. In rodents, one mechanism for dendritic spine formation is through estradiol decreasing GABAergic inhibition in the hippocampus, which in turn increases the excitatory drive on pyramidal cells. The progesterone metabolite 3α,5α-tetrahydroprogesterone (allopregnanolone) blocks this estradiol-induced increase in spine density by altering hippocampal GABAergic activity (58). Allopregnanolone increases tonic inhibition by altering expression of GABA receptor subunits (59–61). Finally, observed effects could stem in part from extra-neuronal factors (62), such as changes in angiogenesis or water content in the brain. In an aerobic exercise intervention study (63) and a recent study of older adults (64), increased hippocampal volume was observed in tandem with increased hippocampal perfusion. Additional studies are needed to determine the extent to which hippocampal volume change is coincident with versus the result of increases in perfusion.

### Clinical implications of the effects of hormone suppression on MTL morphology

Chronic progesterone suppression resulted in robust changes in entorhinal cortex and CA2/3 volume. In response to hormone suppression, CA2/3 volume decreased and entorhinal cortex volume increased relative to naturally cycling conditions, demonstrating hormonal regulation of key structures in the MTL. By far the most pronounced effects of chronic hormone suppression were seen in entorhinal cortex, the first region of the brain to develop neuropathology in the progression of Alzheimer’s Disease (AD) (65). Progesterone acts as an endogenous regulator of β-amyloid metabolism, blocking the beneficial effect of estradiol on β-amyloid accumulation (52). Progesterone also blocks estrogen’s ability to upregulate neurotrophic factors in entorhinal cortex (35). Women are disproportionately affected by AD, a disease in which two-thirds of the patient population are female, and the precipitous decline in sex hormone production during menopause may be a sex-specific AD risk factor (66, 67). Surgical menopause is associated entorhinal cortex thinning (17), and earlier age at menopause (surgical or spontaneous) is associated with an increased risk of cognitive decline and dementia (68–70). Our findings demonstrate strong regulatory effects of progesterone on entorhinal cortex volume, underscoring the need for preclinical and clinical studies to investigate the neuroendocrine basis of cognitive decline and dementia-risk. In a recent study of community-dwelling older adults, high resolution MTL imaging revealed a strong and select correlation between anterolateral ERC volume and performance on the Montreal Cognitive Assessment in individuals at risk for dementia. Few studies of the aging brain consider the midlife period and even fewer consider the neuronal effects of reproductive aging (71). Moving forward, cognitive aging studies should target midlife women to investigate whether changes in progesterone production across the menopausal transition contributes to AD risk, particularly in individuals with genetic susceptibility.

### Further considerations and future studies

The current investigation deeply sampled a single individual across a complete menstrual cycle characterized by canonical fluctuations in sex hormones, and again during chronic progesterone suppression. Though we contend that these results likely reflect general mechanisms of hormonal fluctuations on MTL subregions, future work must enroll a larger sample of women to assess whether individual differences in hormone dynamics influence the relationships observed here.

Next, the present study used peripheral hormone measures to infer a relationship between central hormone levels and brain structure. The hormone regimen (0.02 mg ethinyl-estradiol (EE) and 0.1 mg levonorgestrel (LNG)) used in Study 2 had strong suppressive effects on progesterone, with serum concentrations reduced by ∼97%. Evidence from animal studies indicate that progesterone concentrations in the hippocampus were likely to be blunted as well. In rodents, peripheral steroid hormone levels are correlated with levels in cerebral cortex and hippocampus (72), and a 4-week regimen of the same compound (EE and LNG) suppressed hippocampal concentrations of progesterone (by 65%) and allopregnanolone (by 55%) (73). Exogenous hormone concentrations (conferred by the hormone regimen itself) were also low, with mean EE at 0.01 ng/ml and LNG at 0.91 ng/ml. Despite the overall suppressive effects of the hormone regimen used in Study 2, some degree of progesterone modulation remained, with a 10-fold increase in progesterone between the first and second half of the follow-up study. This residual hormone fluctuation likely underlies the fact that ERC volume retained some degree of modulation in Study 2 (albeit greatly diminished relative to the ERC changes in Study 1).

A long history of clinical evidence implicates hypothalamic-pituitary-gonadal (HPG) axis dysregulation in the development of mood and cognitive disorders (74). Throughout the lifespan, changes in women’s reproductive status have been associated with increased risk for mood disturbance and major depressive disorder (74–80). Given the strong progesterone–MTL volume relationships we report here, it is compelling to speculate about potential consequences on mood and cognition. Though the present analyses were not focused on mood/cognitive changes, the data reported here were collected as part of a larger “28andMe” dataset that included daily assessments of mood/affect and recognition memory (for a detailed description see (81). For completeness, we briefly summarize this data here. Across the cycle, increases in progesterone were associated with increases in negative mood (based on a composite index of the Profile of Mood States; *r* = .37, *p* = .04), with the strongest relationship between progesterone and depression sub-scores across the cycle (*r* = .55, *p* = .001; **Figure S6)**. No associations between sex hormones or MTL volume and cognition were observed, but a cautious interpretation of these findings is warranted given that this was not an *a priori* consideration of the study design (for more, see **Supplementary Results**). Future work should directly investigate whether progesterone-mediated structural changes to the MTL confer changes in mood or cognition. For example, the hippocampus and entorhinal cortex are key regions within the brain’s navigational circuitry, and rodent studies provide powerful evidence that sex hormones play a critical role in spatial navigation ability (82–84). One open question is whether hormone-mediated changes in MTL volume we observed here shapes navigation strategies.

Finally, though the results reported here indicate variability in MTL structure due to hormonal fluctuations across the menstrual cycle, it would be premature to draw the conclusion that variability in MTL structure is greater in females. Data derived from a parallel dense-sampling study in a male (n = 30 days) (85) demonstrate that intra-subject variability in MTL subregion volume does not differ between the sexes. Overall variability was highly similar for both datasets (comparisons were made using unedited data, see Supplementary Results and **Fig. S7**). Further, cross-sectional data from a large population cohort (n=1114; Human Connectome Project Young Adult, S1200 Data Release) show that inter-subject variability in hippocampal volume is similar for males and females (**Fig. S8**). Together, these findings reveal that the variability reported here is on par with the intrinsic variability observed in males. Although we identified a source of variability in subregion volume across the female menstrual cycle (progesterone), this is not an *added* source of variability in females.

While the menstrual cycle is a central facet of the human condition, the female lifespan is punctuated by other periods of hormonal transition (e.g., adolescence, pregnancy, menopause) (71) that also merit deeper interrogation. Applying a dense-sampling, deep-phenotyping lens to these periods in a woman’s life will increase our understanding of brain–hormone interactions significantly.

### Conclusion

Intrinsic fluctuations in progesterone across the menstrual cycle were strongly correlated with CA2/3, entorhinal, perirhinal, and parahippocampal subregion volumes of the human medial temporal lobe. Progesterone suppression abolished these effects. The dynamic endocrine changes that unfold over the menstrual cycle are a natural feature of half the world’s population. Neuroimaging studies that densely sample the individual illuminate the dynamic properties of brain organization. Understanding how endogenous and exogenous changes in sex hormones impact medial temporal lobe morphology is imperative for our basic understanding of the human brain and for women’s health.

## End Notes

### Author contributions

The overall study was conceived by C.M.T., L.P., S.G. and E.G.J.; C.M.T., L.P., T.S., E.L., and E.G.J. performed the experiments; data analysis was conducted by C.M.T., and R.O.; C.M.T. and E.G.J. wrote the manuscript; L.P., R.O., E.L., T.S., and S.T.G. edited the manuscript.

### Data/code availability

MRI data and code will be publicly accessible upon publication.

## Supplementary Information

**Supplementary Figure 1.**
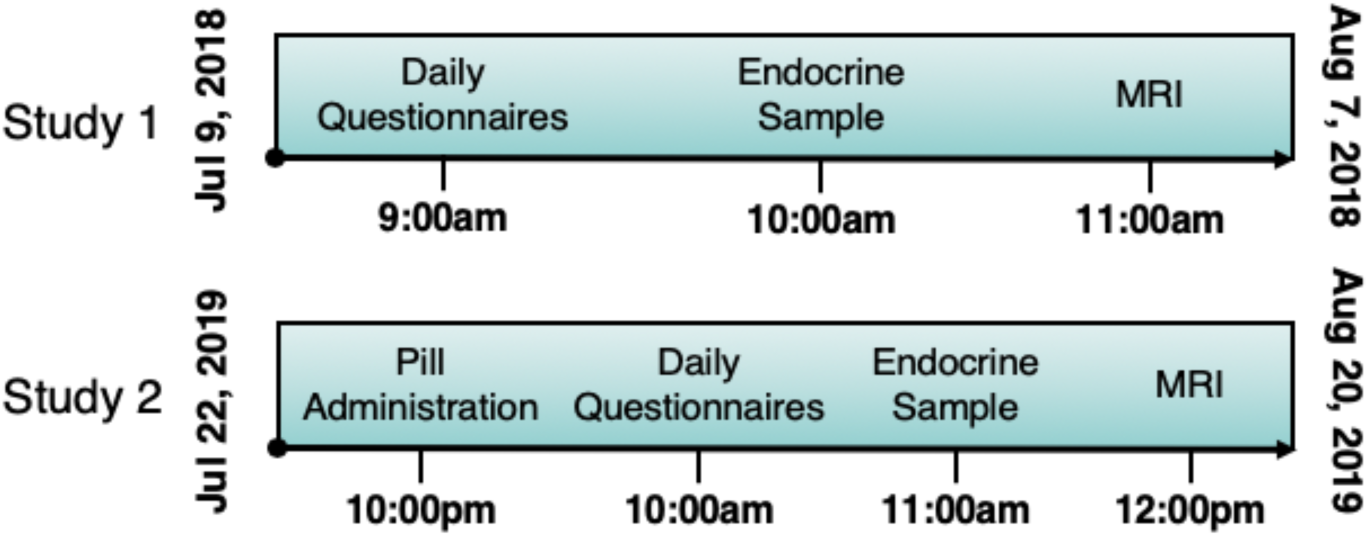
Timeline of data collection for all 60 experimental sessions. Within each study, endocrine and MRI assessments were collected at the same time each day to minimize time-of-day effects. Note that between studies, scan start time was offset by ≤60 minutes due to scheduling constraints. Data collection procedures were identical across the 60 sessions.

**Supplementary Figure 2.**
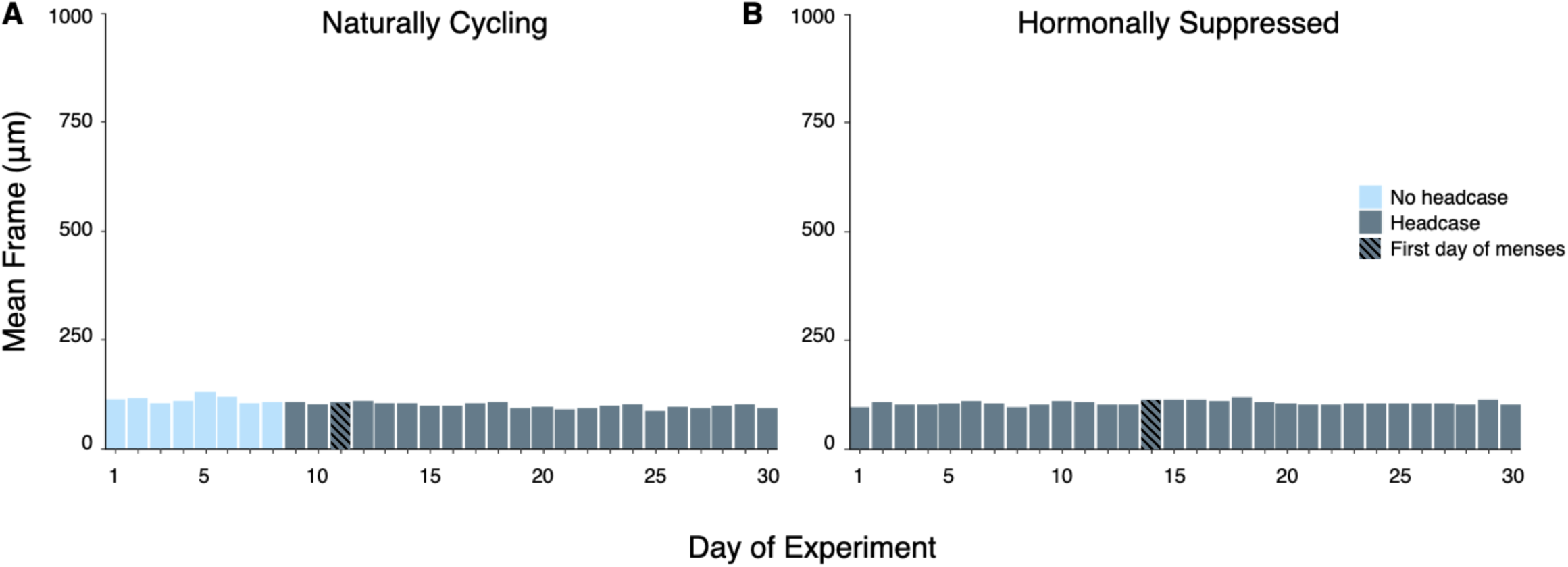
Head motion across the two 30-day experiments. Data reflect motion parameters from a functional scan acquired prior to the hippocampal T2 scan. **A**. During the natural cycle study, motion on days 1–8 was limited using ample head and neck padding. On days 9–30 motion was limited using a molded headcase custom fit to the participant’s head. **B**. Motion parameters (framewise displacement, FWD) during the hormone suppression study. In both studies, movement parameters did not exceed 150 microns.

**Supplementary Figure 3.**
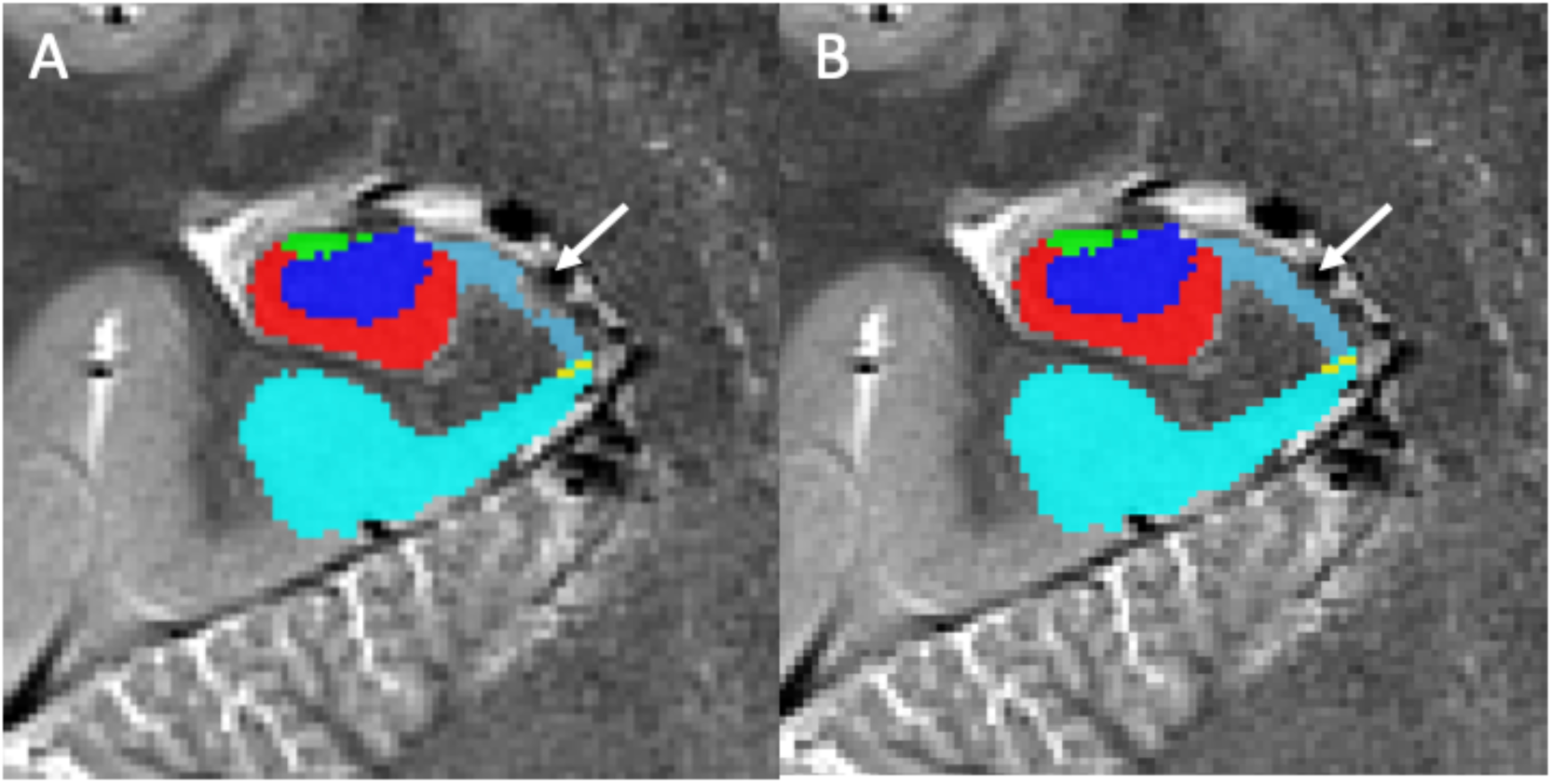
Manual retouching of missing voxels. **A**. Example of original ASHS segmentation with missing/unlabeled voxels within the subiculum. **B**. Segmentation of subiculum after manual retouching.

**Supplementary Figure 4.**
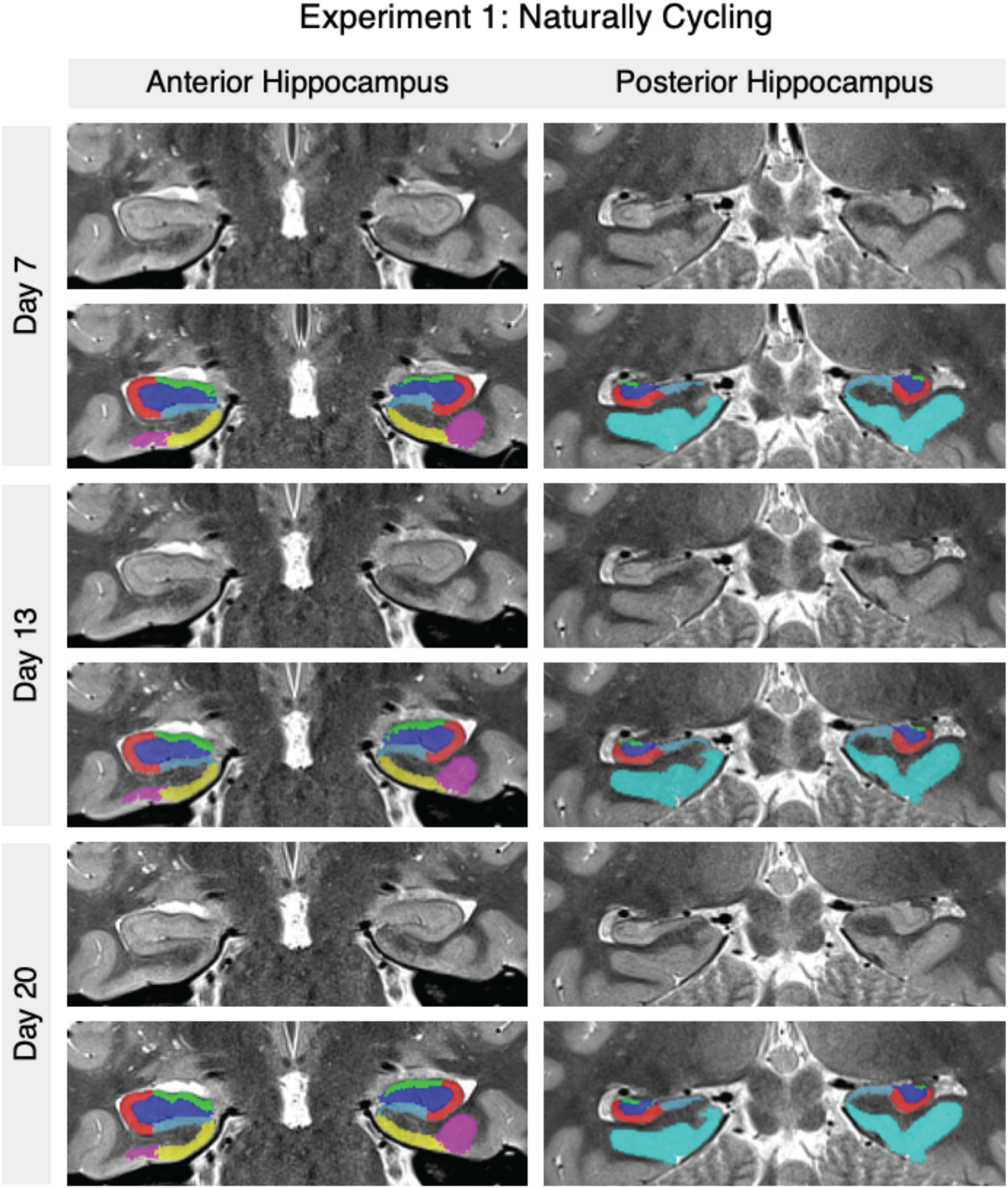
Sample anterior and posterior segmentations. Example of raw images and their segmentations from cycle days 7 and 13 (Phase 1 - low progesterone) and cycle day 20 (Phase 2 - high progesterone).

**Supplementary Figure 5.**
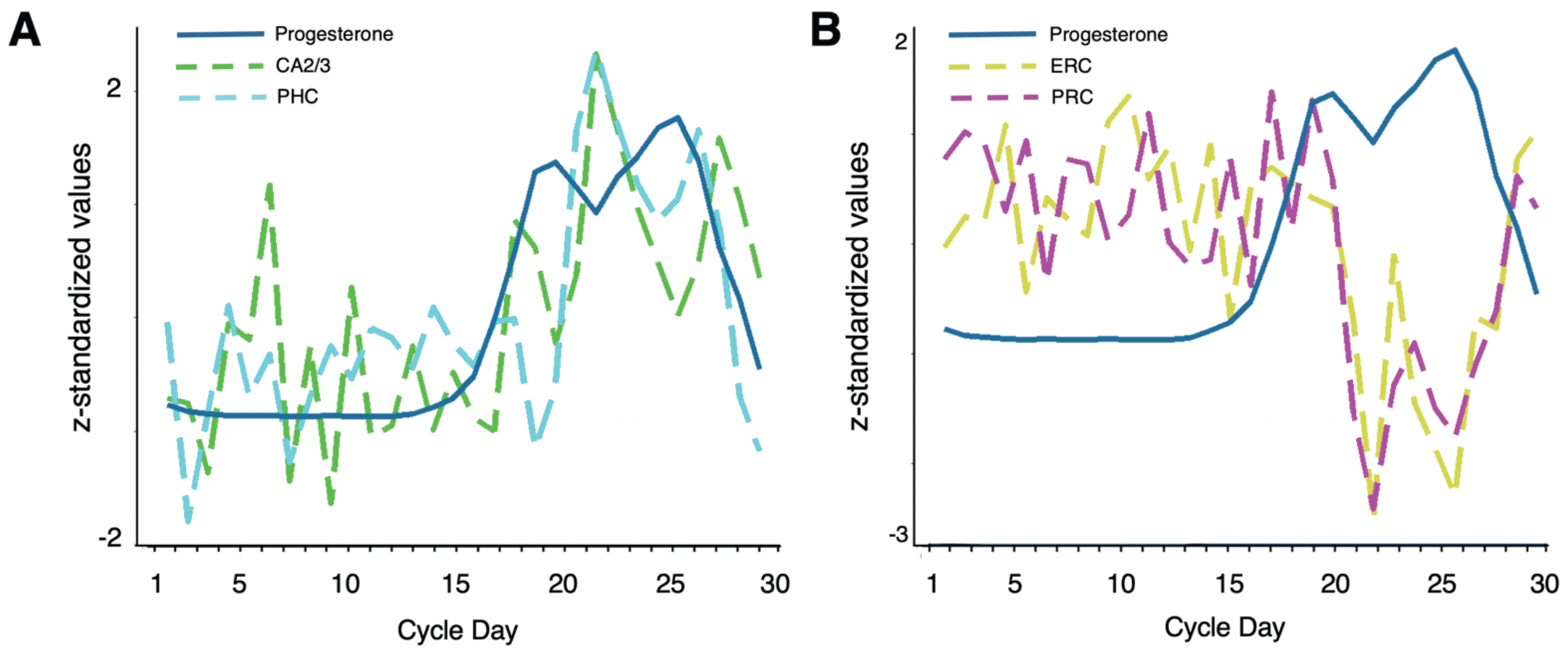
Progesterone and subregion volumes across a menstrual cycle. Time series for standardized progesterone values plotted versus standardized volumes within **A**. positively associated subregions, CA2/3 (R^2^_adj_ = .33, *p* < .001) and parahippocampal cortex (R^2^_adj_ = .38, *p* < .001), and standardized volumes within **B**. negatively associated subregions, entorhinal (R^2^_adj_ = .38, *p* < .001) and perirhinal cortex (R^2^_adj_ = .35, *p* < .001). PHC, parahippocampal cortex; ERC, entorhinal cortex; PRC, perirhinal cortex.

### Behavior across Studies 1 and 2: Supplementary analysis of mood and affect

Data acquisition for Studies 1 and 2 included daily assessments of mood, affect, and recognition memory (for a detailed description see (81). Here we present scores from Study 1 (**Supp. Fig 6A**) and Study 2 (**Supp. Fig 6B**) in relation with progesterone levels. Across the natural cycle, increases in progesterone were associated with increases in negative mood (based on a composite index of the Profile of Mood States; *r* = .37, *p* = .04), with the strongest relationship between progesterone and depression sub-scores (*r* = .55, *p* = .001). No associations between sex hormones or MTL volume and recognition memory were observed.

**Supplementary Figure 6.**
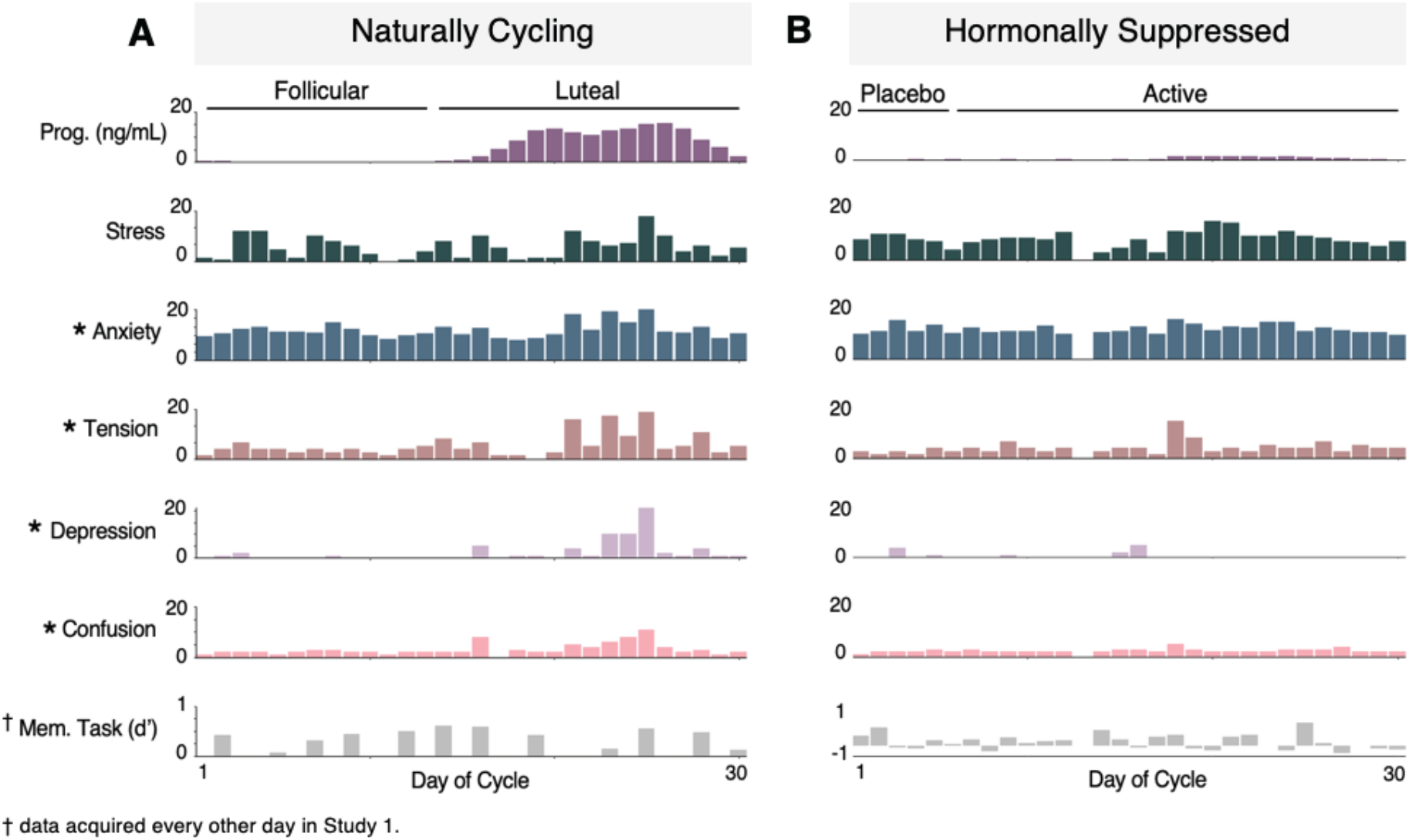
Relationships between progesterone and mood/cognition. Bar plots display daily progesterone and behavioral assessments across both dense-sampling studies. Note that, in Study 1, recognition memory scores were acquired every other day. **A**. Progesterone fluctuations over the cycle were correlated with measures of anxiety (*r*=.36, *p* = .05), tension (*r* = .45, *p* = .013), depression (*r* =.55, *p*=.001), and confusion (*r* = .52, p = .003). **B**. Hormone-mood associations were not observed within the hormone suppression study, however daily depression scores were marginally lower during chronic hormone suppression compared to freely cycling (*p* = .044) while self-reported stress scores were higher (*p* = .008). *\** denotes *p* ≤ .05

**Table S1.** Gonadal and pituitary hormones by menstrual cycle phase and hormonal contraceptive phase

|  | **Estradiol**<br>**(pg/l)** | **Progesterone**<br>**(ng/l)** | **LH**<br>**(mIU/mL)** | **FSH**<br>**(mIU/mL)** |
| --- | --- | --- | --- | --- |
| **Follicular** |  |  |  |  |
| Mean (SD) | 37.9 (15.9) | 0.2 (0.2) | 5.9 (0.7) | 6.5 (1.2) |
| Standard range | 12.5-166.0 | 0.1-0.9 | 2.4-12.6 | 3.5-12.5 |
| **Ovulatory** |  |  |  |  |
| Mean (SD) | 185.3 (59.0) | 0.2 (0.2) | 21.7 (16.4) | 8.1 (3.6) |
| Standard range | 85.8-498.0 | 0.1-12.0 | 14.0-95.6 | 4.7-21.5 |
| **Luteal** |  |  |  |  |
| Mean (SD) | 85.4 (26.4) | 9.5 (4.8) | 5.5 (2.0) | 4.8 (1.3) |
| Standard range | 43.8-210.0 | 1.8-23.9 | 1.0-11.4 | 1.7-7.7 |
| **Placebo** |  |  |  |  |
| Mean (SD) | 21.3 (10.3) | 0.06 (0.01) | 6.1 (1.4) | 7.6 (1.0) |
| **Active** |  |  |  |  |
| Mean (SD) | 79.9 (77.6) | 0.18 (0.13) | 6.3 (3.6) | 4.2 (1.2) |

Note: Standard reference ranges based on aggregate data from Labcorp (https://www.labcorp.com)

**Table S2.** Volumes (mm^3^) and statistics of hippocampal subfields by hemisphere – Naturally cycling

|  | **Total Hipp** | **CA1** | **CA2/3** | **DG** | **Sub** | **ERC** | **PRC** | **PHC** |
| --- | --- | --- | --- | --- | --- | --- | --- | --- |
| **Left Hemisphere** |  |  |  |  |  |  |  |  |
| **Average**<br>(SD) | 2805.6<br>(42.2) | 647.0<br>(17.6) | 154.8<br>(16.2) | 722.3<br>(34.5) | 466.2<br>(22.3) | 617.1<br>(33.1) | 1621.1<br>(196.6) | 2807.7<br>(176.5) |
| **Min** | 2734.4 | 618.3 | 129.1 | 657.3 | 398.9 | 529.8 | 1164.6 | 2503.4 |
| **Max** | 2888.8 | 681.8 | 195.0 | 780.3 | 487.7 | 672.3 | 1904.0 | 3191.2 |
| **Range**<br>(% of Max Volume) | 154.4<br>(5.3) | 63.5<br>(9.3) | 65.9<br>(33.8) | 123.0<br>(15.8) | 88.8<br>(18.2) | 142.5<br>(21.2) | 739.4<br>(38.8) | 687.9<br>(21.6) |
| **Correlation with E**<br>*p*-value (*r*) | n.s. | n.s. | n.s. | n.s. | n.s. | n.s. | n.s. | n.s. |
| **Correlation with P**<br>*p*-value (*r*) | n.s. | 0.0125<br>(0.45) | 0.0202‡<br>(0.42) | 0.0040†<br>(0.51) | n.s. | 0.0028*†<br>(-0.53) | 0.0002*†<br>(-0.63) | 0.0003*†‡<br>(0.61) |
|  | **Total Hipp** | **CA1** | **CA2/3** | **DG** | **Sub** | **ERC** | **PRC** | **PHC** |
| **Right Hemisphere** |  |  |  |  |  |  |  |  |
| **Average**<br>(SD) | 3100.9<br>(54.6) | 700.0<br>(18.4) | 248.5<br>(23.0) | 726.5<br>(20.9) | 424.8<br>(21.4) | 775.8<br>(46.5) | 1073.7<br>(106.7) | 2516.7<br>(147.8) |
| **Min** | 2952.0 | 647.0 | 201.7 | 676.6 | 379.6 | 644.2 | 850.2 | 2265.0 |
| **Max** | 3202.8 | 741.9 | 310.1 | 780.3 | 463.3 | 836.5 | 1223.8 | 2862.9 |
| **Range**<br>(% of Max Volume) | 250.9<br>(7.8) | 94.9<br>(12.8) | 108.3<br>(34.9) | 103.7<br>(13.3) | 83.7<br>(18.1) | 192.3<br>(23.0) | 373.5<br>(30.5) | 597.8<br>(20.9) |
| **Correlation with E**<br>*p*-value (*r*) | n.s. | n.s. | n.s. | n.s. | n.s. | n.s. | n.s. | n.s. |
| **Correlation with P**<br>*p*-value (*r*) | n.s. | 0.0248<br>(-0.41) | 0.0125‡<br>(0.45) | n.s. | n.s. | 0.0003*†<br>(-0.61) | 0.0166<br>(-0.43) | 0.0005*†‡<br>(0.60) |

n.s. indicates $p > 0.05$ \* indicates significance at Bonferroni corrected $p < 0.0036$ † indicates significance at $p < 0.05$ with GMV included in linear model‡ indicates significance at $p < 0.05$ with slice number included in linear model

**Table S3.** Volumes (mm^3^) and statistics of hippocampal subfields by hemisphere – Hormonally suppressed

|  | CA1 | CA2/3 | DG | Sub | ERC | PRC | PHC |
| --- | --- | --- | --- | --- | --- | --- | --- |
| **Left Hemisphere** |  |  |  |  |  |  |  |
| **Average**<br>(SD) | 627.9<br>(15.7) | 133.9<br>(11.0) | 705.4<br>(27.9) | 433.0<br>(22.0) | 688.0<br>(18.4) | 1648.2<br>(166.8) | 2746.9<br>(109.5) |
| **Min** | 595.7 | 118.1 | 666.5 | 397.0 | 650.0 | 1381.2 | 2496.0 |
| **Max** | 658.9 | 164.2 | 774.8 | 484.6 | 727.5 | 2086.5 | 2920.5 |
| **Range**<br>(% of Max Volume) | 63.2<br>(9.6) | 46.1<br>(28.1) | 108.3<br>(14.0) | 87.6<br>(18.1) | 77.5<br>(10.7) | 705.3<br>(33.8) | 424.5<br>(14.5) |
| **Correlation**<br>**with E**<br>*p*-value (*r*) | n.s. | n.s. | n.s. | n.s. | 0.0257<br>(-0.41) | 0.0140<br>(0.45) | n.s. |
| **Correlation**<br>**with P**<br>*p*-value (*r*) | n.s. | n.s. | n.s. | n.s. | 0.0107<br>(-0.47) | n.s. | n.s. |
|  | CA1 | CA2/3 | DG | Sub | ERC | PRC | PHC |
| **Right Hemisphere** |  |  |  |  |  |  |  |
| **Average**<br>(SD) | 722.9<br>(31.1) | 223.6<br>(15.8) | 728.0<br>(30.2) | 423.3<br>(22.7) | 986.2<br>(26.9) | 1051.1<br>(111.0) | 2489.8<br>(102.7) |
| **Min** | 688.2 | 181.0 | 683.9 | 388.8 | 793.5 | 952.5 | 2305.0 |
| **Max** | 829.8 | 254.5 | 803.5 | 505.1 | 922.5 | 1487.7 | 2637.6 |
| **Range**<br>(% of Max Volume) | 141.6<br>(17.1) | 73.5<br>(28.9) | 119.6<br>(14.9) | 116.3<br>(23.0) | 129.1<br>(14.0) | 535.3<br>(36.0) | 332.6<br>(12.6) |
| **Correlation**<br>**with E**<br>*p*-value (*r*) | n.s. | 0.0474<br>(-0.37) | n.s. | n.s. | n.s. | n.s. | n.s. |
| **Correlation**<br>**with P**<br>*p*-value (*r*) | n.s. | n.s. | 0.0035*<br>(-0.52) | n.s. | n.s. | n.s. | n.s. |

Note: Cycle Day 13 volumes were considered outliers and not included in summary statistics or analyses
n.s. indicates $p > 0.05$
\* indicates significance at Bonferroni corrected $p < 0.0036$

### Comparison of MTL variability by sex: Supplementary analysis of dense-sampling data in a male

#### Background

In 2011, Beery and Zucker (1) established the striking sex bias in biomedical research. Across most biomedical scientific disciplines, neuroscience chief among them, females are disproportionately excluded from preclinical research. One foundational argument for this exclusion is that the estrous cycle renders females too variable to include in research. This assumption failed empirical testing. Prendergast et al. (2) issued a report in mice showing that across 9,932 trait measurements taken from 293 independent articles, unstaged females were no more variable than males in any trait, and males showed higher overall trait variability.

Though males and females are largely recruited in equal numbers in human brain imaging studies, the assumption of greater variability in females is still being used as justification for excluding females from certain investigations. To address this point head-on, we capitalized on existing data from two sources: a dense-sampling study in a male for which high resolution MTL imaging was available over a similar time-course (n = 30 days); and a large-scale dataset assessing total hippocampal volume in men and women.

Data from the densely-sampled male were used to determine the magnitude of total *intra*-subject variability in MTL volume in the male relative to our densely-sampled female. Data from the population-based study were used to assess *inter*-subject variability by sex. Comparisons of variance confirmed few differences in overall intra-subject variability between sexes and greater inter-subject variability in males. Details of the companion datasets and supplementary analyses are provided below.

#### Determining Intra-Subject Variability by Sex

To determine whether variability in MTL subregion volume across the menstrual cycle reflected intra-subject variability unique to females, we compared the variability of MTL subregion volumes reported here to that of a densely-sampled male.

##### Male “Day2Day” Dataset

Data from a male subject were acquired through the publicly available Day2Day dataset (3). For this dataset, eight participants (2 male, mean age 29 years, SD = 2.58, range 24–32) underwent MRI scanning on multiple (40-50) occasions over the course of 6-8 months. For this analysis, data from study subject 07 (male) was selected, since the participant had been scanned at least 30 times (n = 43 scans). In order to match the session size with the female data reported in the main text, data from the first 30 scans were used.

##### MRI Acquisition

Scanning parameters are described in (3), and were similar to those reported here.

##### Hippocampal Segmentation

*T*_*1*_- and *T*_*2*_-weighted scans were submitted to the automatic segmentation of hippocampal subfields package (ASHS) as described in the main text. *T*_*2*_ scans and segmentations were visually examined using ITK-SNAP for quality assurance. No manual editing was conducted on the male segmentation data.

##### Statistical Analysis

All analyses were conducted in R (version 3.4.4). As no manual editing was conducted on the male data, statistical comparisons were conducted with unedited volumes from our dataset. To compare MTL subregion variance between our female and male data, we conducted F-tests on volumes within each MTL subregion. There was no significant difference in variance observed in CA1 (F_(1,29)_ = 0.828, *p* = .62), DG (F_(1,29)_ = .485, *p* = .06), Sub (F_(1,29)_ = 1.59, *p* = .22), PRC (F_(1,29)_ = 1.95, *p* = .08) or PHC (F_(1,29)_ = .524, *p* = .09). There were significant differences observed in CA2/3 (F_(1,29)_ = 0.450, *p* = .04; M > F) and ERC (F_(1,29)_ = 2.84, *p* < .01; F > M). For visualization of these comparisons, subregion volumes were mean-centered and plotted in R. (**Supp. Fig. 7**).

**Supplementary Figure 7.**
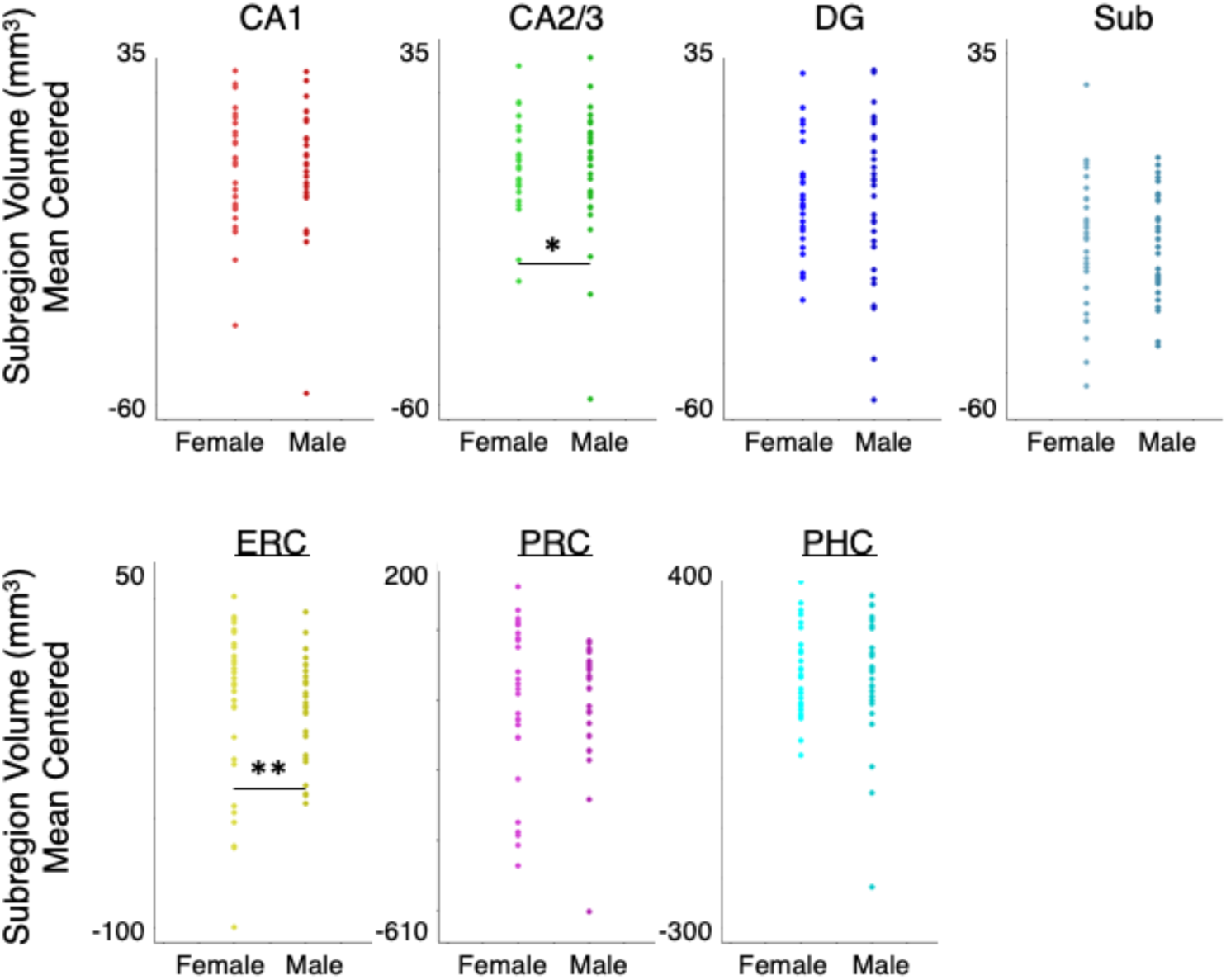
Intra-subject variability in MTL subregions volume by sex. Data depict female and male mean-centered MTL subregion volumes. Data derived from a dense-sampling study in a male exhibit similar *intra-*subject variability in MTL volume compared to our densely-sampled female. Variance was highly similar for 5 of the 7 subregions. Greater variance in female subregion volume was observed in ERC, and greater variance in males was observed in PHC. \**p* < .05, \*\**p* < .01

#### Determining Inter-Subject Variability by Sex

To determine whether variability in hippocampal volume differs significantly between males and females, we compared variance in total hippocampal volume using a large, population-based sample.

##### Human Connectome Project Data

Data were acquired through the publicly available Human Connectome Project (4) Young Adult dataset (S1200 Data Release, n = 508 males, 606 females with imaging data; aged 22–35). For this analysis, preprocessed structural data were used (5), specifically whole hippocampal volumes (6).

##### Statistical Analysis

All analyses were conducted in R (version 3.4.4). To account for differences in brain size between males and females, participants’ hippocampal volumes (average of left and right) were divided by total brain volume (sum of total gray and white matter volumes). An F-test was performed to compare variance in TBV-corrected hippocampal volumes between males and females. There was no difference in inter-subject variability in hippocampal volume between males and females (F_(1,605)_ = 1.04, *p* = .65). For visualization of this comparison, TBV-corrected hippocampal volumes were mean-centered and plotted in R. (**Supp. Fig. 8**).

**Supplementary Figure 8.**
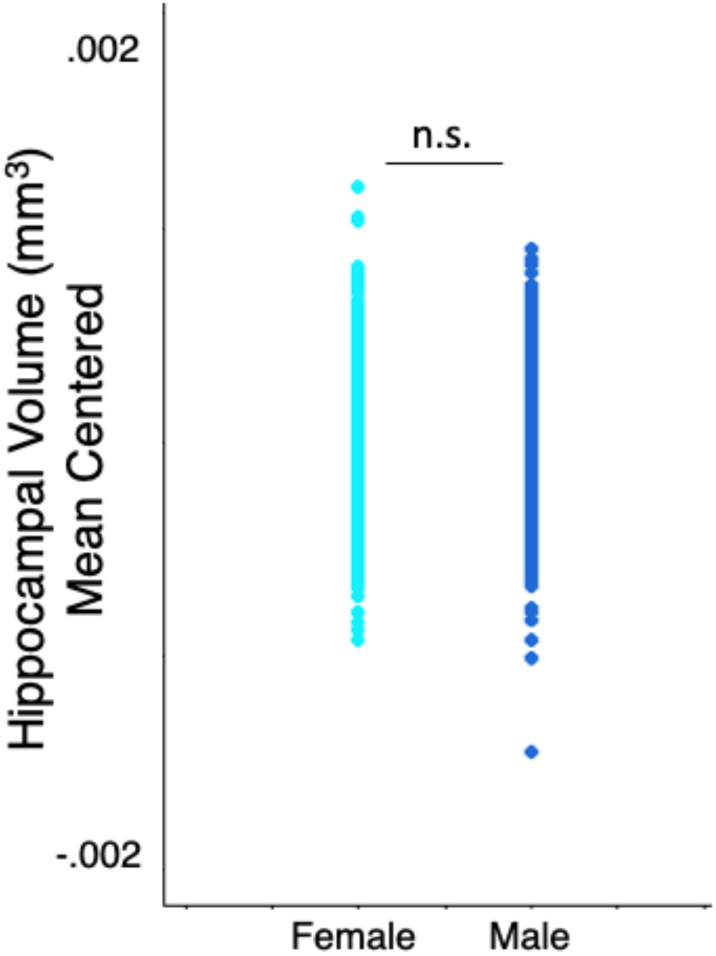
Inter-subject variability in hippocampal volume by sex. Cross-sectional data from a large population cohort (n=1114; Human Connectome Project Young Adult, S1200 Data Release) show that inter-subject variability in hippocampal volume (adjusted for total brain volume) is not different between males and females.

#### Conclusions

In summary, overall intrasubject variability in MTL subregion volume is largely similar between sexes. Further, data derived from a large population cohort (n = 1113) indicate that intersubject variability in hippocampal volume may be larger in males than females. Together, these findings indicate that fluctuations in progesterone shape MTL volume over time, however this variability is on par with the variability observed in males.

## Acknowledgements

We thank Simone Kühn for access to the Day2Day dataset and the Human Connectome Project for access to their population-based dataset.

This work was supported by the Harvey L. Karp Discovery Award (CMT), the Brain and Behavior Research Foundation (EGJ), the California Nanosystems Institute (EGJ), the Hellman Family Fund (EGJ) and the Rutherford B. Fett Fund (STG). Thanks to Mario Mendoza for phlebotomy and MRI assistance. We would also like to thank Shuying Yu, Courtney Kenyon, Maggie Hayes, and Morgan Fitzgerald for assistance with data collection.

